# Permeation of cisplatin through the membranes of normal and cancer cells: a molecular dynamics study

**DOI:** 10.1101/375980

**Authors:** T. Rivel, C. Ramseyer, S. O. Yesylevskyy

## Abstract

In this work, realistic models of membranes of normal and cancer cells are developed. A special focus is given to their cholesterol content. It is shown that the loss of lipid asymmetry in the membranes of cancer cells leads to a decrease of their permeability to cisplatin by one order of magnitude in comparison to the membranes of normal cells. The change of cholesterol molar ratio from 0% to 33% also decreases the permeability of the membrane by approximately one order of magnitude. The permeability of pure DOPC membrane is 5-6 orders of magnitude higher than one of the membrane with realistic lipid composition, which makes it as an inadequate model for the studies of drug permeability.

## Introduction

Although the structure and major properties of the cell membranes are considered to be known since the introduction of the fluid-mosaic model (1), there are still many open questions concerning membrane composition, permeability and mechanical properties. Continuous increase of computer power stimulated rapid development of computational approaches to the modeling of realistic cell membranes using both atomistic and coarse-grained molecular dynamics (MD) techniques (2-8). Particularly, recent works address such phenomena as lateral heterogeneity of the membranes (lipid rafts and micro-domains, lipid sorting) (2, 3); asymmetry of the lipid composition and cholesterol content between the monolayers (9); membrane curvature and its influence on the physical properties of bilayer (10) and so on.

The existence of lipid rafts or micro/nano-domains in the membranes is still a debated concept. It is shown that cholesterol (CHL) and sphingomyelin lipids (SM) are forming regions of rigid liquid-ordered phase (Lo) *in vitro* in giant unilamellar vesicles (GUVs), in liposomes or in deposited lipid structures (11-13). The rafts were also extensively studied *in silico* (11, 14-16). However, there are difficulties in observing the rafts in real cells, which are usually attributed to their small life time or to their size, which appears to be much smaller in real cells in comparison to artificial membranes (14, 17, 18). Despite the fact that the rafts are not observed directly in living cells, the influence of cholesterol and cholesterol-rich microdomains on the membrane rigidity, on permeability for small molecules and on the functioning of the membrane proteins are of great interest (19-22).

An asymmetry in the lipid composition between the monolayers of the plasma membrane is now recognized as an important factor of membrane functioning. The composition of the membrane leaflets is highly uneven (13, 23, 24) and is actively maintained by a group of proteins – the flippases and floppases. This results in an extracellular leaflet composed mostly of phosphatidylcholine (PC) and SM and a cytoplasmic leaflet enriched in phosphatidylserine (PS) and phosphatidylethanolamine (PE). The transversal distribution of CHL is subject to debates. It has been observed experimentally (25-27) that CHL resides mostly in the cytoplasmic leaflet. On the other hand, the affinity of cholesterol to SM has been shown experimentally (18) and *in silico* simulations also tend to show an increased distribution of CHL to the outer leaflet (9, 28, 29). Although more and more computational studies are trying to mimic an actual asymmetric lipid content of the membrane monolayers (4, 5, 7, 10), we are not aware of any works which are addressing the question of the influence of this asymmetry on the permeability of the membranes to drugs and small molecules.

Besides the asymmetry of the membrane, its curvature also plays an important role in many cellular processes (26, 30). The curvature was recently shown to induce changes in the distribution of cholesterol in the membrane, in the thickness of its leaflets and in the order parameter of the lipid tails (10) but the studies of the influence of curvature on the passive diffusion of drugs and small molecules through the membrane are scarce.

Accounting for membrane asymmetry and realistic lipid composition in MD simulations is especially important due to the fact that this composition changes significantly in malignant cells. Since 1989 it is known that some cancer cells expose PS on the extracellular leaflet of their plasma membranes, while this anionic lipid is predominantly located in the intracellular leaflet in the normal cells (31, 32). Subsequent studies using flow cytometry after labeling by annexin V confirmed these findings (33-35). It was also shown that exposure of PS in cancer cells is not an artifact caused by the presence of apoptotic cells in the sample (36). PS exposure was also detected in the vasculature of the tumors (5, 37). Table S1 in Supplementary Information summarizes existing literature about PS exposure in different cancers and cancer cell lines. It is evident that the outer monolayer of cancer cells contains on average ~7 times more PS than the normal control cells. These findings resulted in a new approach in targeting tumors by selective recognition of exposed PS (35, 38-41) or related redistribution of PE (42). Despite growing importance of this field we have found only one work where *in silico* modeling of the membranes of cancer cells was performed (6).

MD simulations provide a unique opportunity of studying passive diffusion of drugs and small molecules through realistic model membranes. All-atom MD simulations are able to provide accurate quantitative results on permeability coefficients and diffusion constants of the studied compounds in the membranes. Permeability is usually estimated using inhomogeneous solubility-diffusion (ISD) formalism (43, 44), which accounts for transversal heterogeneity of the membrane along its normal. The ISD model was used successfully in many studies (45, 46) (see (47) for extensive review). However, a number of publications report consistent difficulties in computing free energy profiles and diffusion coefficients and discuss possible improvements in ISD methodology (48-52).

Permeability of the membranes of normal and cancer cells of the anti-tumor drugs, which are currently available on the market or in the clinical studies pipeline, is of great interest. It is obvious, however, that some well-known and widely used drug should be studied first to show that the difference in permeability between normal and cancer cell membranes exists. Cisplatin (*cis*-diamminedichloroplatinum(II)) is a good candidate for such study. It is one of the oldest and the most widely used anti-tumor agents, which is adopted for treatment of a wide range of cancers (53). While its action, which leads to cell death by apoptosis or necrosis by means of DNA damage and redox stress, is well studied (see (54, 55) for reviews), the mechanism of its transport trough the membrane is not yet understood. There were numerous studies (56-60) which supported the hypothesis that the copper transporter CTr1 facilitates the transport of cisplatin into the cell. However, more recent works mostly exclude this hypothesis (53, 61-64). Although some studies show a positive correlation between the presence of CTr1 in the membrane and the high transport rate of cisplatin, they do not necessarily show an increase of the cell death rate. This may suggest that cisplatin appears in the cytoplasm in inactivated chemical form after being transported by CTr1 (61, 65-67). Currently passive diffusion through the lipid bilayer is considered as the only known way of transport of cisplatin in its active form (53, 58, 65, 68).

The chemistry of solvated cisplatin is complex (69). It undergoes numerous reactions such as oligomerization, hydrolysis, hydroxylation, protonation/deprotonation, substitution of the chloride with carbonate ions, sulfur groups or other nucleophile moieties, *etc*. (70-82). It is shown that the native form of cisplatin has the highest propensity to passively diffuse through the plasma membrane and is the most abundant form in the extracellular medium (83-85).

It is possible to conclude that atomistic models of cell membranes with realistic lipid composition and cholesterol distribution are of high demand for both normal and cancer cells. The computations of permeability of such membranes for drugs are of great importance for on-going attempts of targeting the cancer cells by their membrane composition. Moreover, the methodology of computing the permeability coefficients is well-established and tested. Despite this interest there are no dedicated works which are focused on computing permeabilities in normal and cancer cell membranes for common anti-tumor drugs. There are also very few systematic attempts to compare permeabilities of the membranes with different cholesterol content in complex model membranes.

In this work we computed permeabilities of realistic model membranes of normal and cancer cells for widely used anti-cancer drug cisplatin. We also compared permeabilities in the membranes with different cholesterol content ranging from 0% to 33% molar ratio. It is shown that the cancer cell membrane is approximately one order of magnitude less permeable to cisplatin in comparison to the membrane of the normal cell. We discuss possible reasons of this difference in details and compare obtained results with available experimental data.

## Methods

### Model membranes

We utilized the model of realistic plasma membrane, referred as “normal membrane” hereafter, developed in our previous work (10). In this model the asymmetry of real mammalian plasma membranes is taken into account – the outer monolayer is enriched in sphingomyelin (SM) and 1,2-dioleoyl-sn-glycero-3-phosphocholine (DOPC), while the inner monolayer is enriched in 1,2-dioleoyl-sn-glycero-3-phosphoethanolamine (DOPE) and 1,2-dioleoyl-sn-glycero-3-phospho-L-serine (DOPS).

The model of the plasma membrane of a cancer cell, referred as “cancer membrane” hereafter, was built using the same protocol as described in (10) but with increased proportion of DOPS and DOPE in the extracellular leaflet. Figure 1 shows a schematic representation of the normal and cancer membranes along with their lipid compositions and comparison to available experimental data. Table 1 shows the lipid content of the monolayers of normal and cancer membranes. The normal membrane was designed according to well-established lipid content of mammalian erythrocyte membrane (86). In the case of the cancer membrane the distribution of the lipid species was symmetrized to emphasize the overexpression in PS/PE in the extracellular leaflet. The Slipids force field (87) was used for lipids, which is one of the best force fields for the membrane systems available today (88).

**Table 1.** Lipid content (absolute number of molecules) of the monolayers of simulated membranes.

| Component | Normal |  | Cancer |  |
| --- | --- | --- | --- | --- |
|  | Outer | Inner | Outer | Inner |
| SM (sphingomyelin) | 42 | 12 | 27 | 27 |
| PC (1,2-dioleoyl-sn-glycero-3-phosphocholine) | 46 | 14 | 30 | 30 |
| PE (1,2-dioleoyl-sn-glycero-3-phosphoethanolamine) | 14 | 46 | 30 | 30 |
| PS (1,2-dioleoyl-sn-glycero-3-phospho-L-serine) | 0 | 30 | 15 | 15 |
| Cholesterol | 51 | 51 | 51 | 51 |

**Figure 1.**
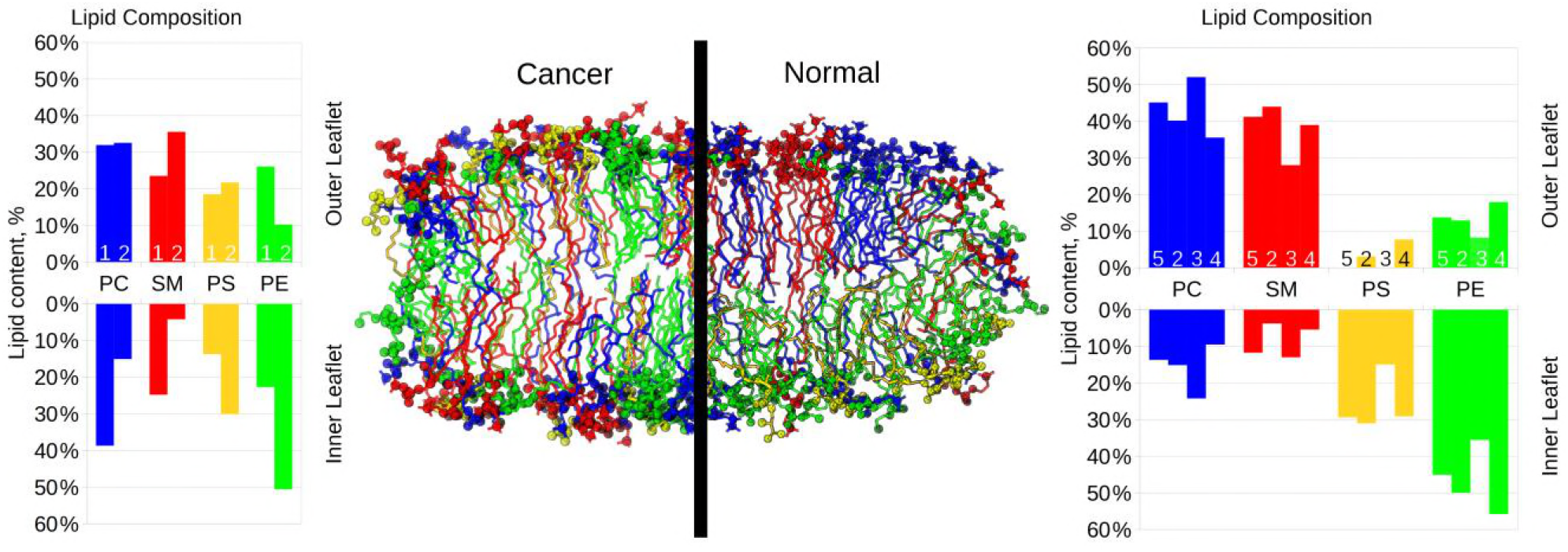
Snapshots of the simulated systems for cancer and normal membrane models. DOPC is shown in blue, SM in red, DOPS in yellow and DOPE in green. For the sake of clarity cholesterol is not shown. Head groups of lipids are shown as spheres. The histograms show the relative abundance of different lipid species for inner and outer monolayers for each membrane model (normalized for each monolayer). Numbers 1 and 5 correspond to the cancer and normal models presented in this work respectively, model 2 refers to the work by Klähn and Zacharias (6), model 3 to the work by Ingólfsson *et al*. (4), model 4 to the data reported by Marquardt *et al*. (26).

The membrane was prepared as a bicelle which is limited by cylindrical caps in XZ plain and forms an infinite bilayer in Y direction. This allows the monolayers to relax to their optimal areas by exchanging the lipids with the bicelle caps, which serve as “compensatory reservoirs”. The mixing of the lipids from different monolayer was prevented by artificial repulsive potentials as described in our previous works (9, 89, 90). This setup was recently used with great success for both atomistic and coarse-grained studies of curved membranes. The reader is referred to (10, 89) for detailed rationale and discussion of this technique.

Initial bicelle has a cholesterol molar content of 33% relative to the lipids. We also constructed the systems with cholesterol content of 0% and 15%. The later was made by randomly removing half of cholesterol molecules and the former by removing all of them. An equal amount of cholesterol was removed from both monolayers in the case of 15% content.

The reference single-component DOPC membrane was also simulated. This system was also arranged as a bicelle using the same setup except the fact that no repulsive potential is applied in the caps since the mixing of the monolayers is allowed in this case.

All membranes were equilibrated for at least 200 ns before computing the permeabilities.

### Computations of permeability

The QDsol topology of cisplatin from our previous work was used (91). This topology reproduces internal flexibility of the coordination bonds of platinum atom accurately by fitting them to the values obtained in *ab initio* quantum dynamics of solvated cisplatin. This model is certainly the best topology of cisplatin available today.

The potentials of mean force (PMF) of translocation of cisplatin were computed using the umbrella sampling method. Three to five cisplatin molecules were placed in equal intervals along the X axis at the bilayer part of the bicelle. The harmonic biasing potential with the force constant of 1000 kJ/mol/nm^2^ was applied to the centre of mass of each ligand in Z dimension, which is perpendicular to the membrane plane. The potentials were centered at discrete points distributed along Z axis in 0.1 nm intervals. Additional weak flat-bottom potential was applied along X axis for each ligand to prevent their accidental interaction due to uncontrollable lateral diffusion. This produced 50 to 57 umbrella sampling windows spanning through the whole bilayer along Z axis. The PMFs were computed as average of all ligands using the weighted histogram technique as implemented in Gromacs. Each window was simulated for at least 16 ns with the last 5 ns used for sampling for a total of 6.25 μs. A plot showing the convergence of the free energy profile is reported in the supplementary information (Fig. S1).

The diffusion coefficients D of the ligands were computed for each umbrella sampling window using the force correlation method (92):

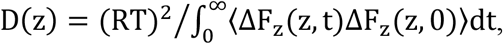

where *R* is the gas constant, *T* is an absolute temperature, ΔF_z_(z, t) is the difference between the instantaneous force and mean force acting on the permeant at given time *t*. The autocorrelation functions very fitted by single exponential decay prior to integration as implemented in Gromacs tools. Diffusion coefficients of all the ligands were averaged for each window. In order to verify the validity of calculations the diffusion coefficient was also computed using the mean-square displacement of the ligands (see Supplementary Information).

The permeability coefficients *P* of cisplatin were computed using standard inhomogeneous solubility-diffusion model (92):

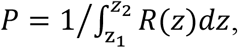

where *R*(*z*) is the local permeation resistance of the membrane at depth *z* expressed as

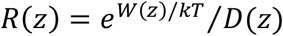

where *W*(*z*) is the PMF of a given ligand. The integration limits *z*_1_ and *z*_2_ were chosen as ±2.3 nm from the membrane center (the points in the water phase outside the membrane). The value of *P* is insensitive to the choice of these limits since the resistance *R* in water phase is negligible.

### Technical details

All MD simulations were performed in Gromacs (93) versions 5.1.2 and 2016.1 in NPT ensemble at the pressure of 1 atm maintained by Berendsen barostat (94) with anisotropic pressure coupling along Y direction only (see (10, 89) for rationale of this choice). The cutoff for non-bonded interactions was 0.8 nm with Verlet cutoff scheme (95). Long range electrostatics was computed with the PME method (96). Velocity rescale thermostat (97) was used. The temperature of 320K was used. An integration steps of 1 fs was used in all simulations as required by the “soft” bonds of cisplatin topology (10). No bonds were converted to rigid constraints.

## Results

### Properties of the membranes

It is well known that cholesterol content changes physical properties of the membrane significantly, in a way referred in the literature as the “condensing effect|” (19). In order to assess these changes in our systems we computed the density profiles of head groups and tails (Fig. 2) and the order parameter of the lipid tails (Fig. 4.). These observables were averaged over the last 10 ns of each umbrella sampling windows used for the computation of the PMFs of cisplatin, which sums up to 0.5 μs of total simulation time.

**Figure 2.**
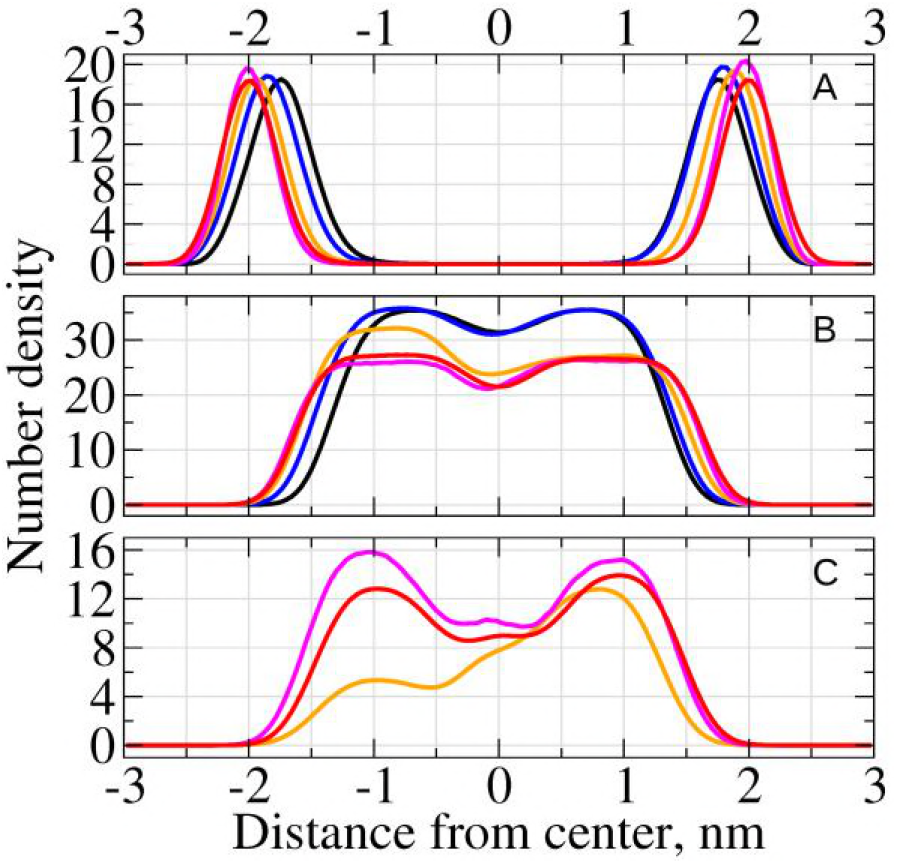
Number densities of A. the lipid head groups, B. the lipid tails and C. the cholesterol for different model membranes. Outer leaflet corresponds to positive distances from the membrane center. The following color convention was used in all figures unless stated otherwise: DOPC model is in black, normal model (0% CHL) is in blue, normal model (15% CHL) is in yellow, normal model (33% CHL) is in fuchsia, cancer model (33% CHL) is in red.

It is clearly visible from Fig. 2 that the distance between the head groups (the thickness of the membrane) increases with increase of cholesterol content. This trend is shown as a function of cholesterol content in Fig. 3. The trends for the inner and the outer leaflets of the normal membrane exhibit slightly different slopes but both of them demonstrate perfect linear dependence (the correlation coefficients of linear regression are 0.998 and 0.988 respectively). The pure DOPC bilayer is the thinnest, which is expectable taking into account a more compact conformation of its unsaturated lipid tails. The cancer membrane with 33% cholesterol has identical thicknesses of both monolayers which is comparable with the thickness of the outer leaflet of the normal membrane with the same cholesterol content.

**Figure 3.**
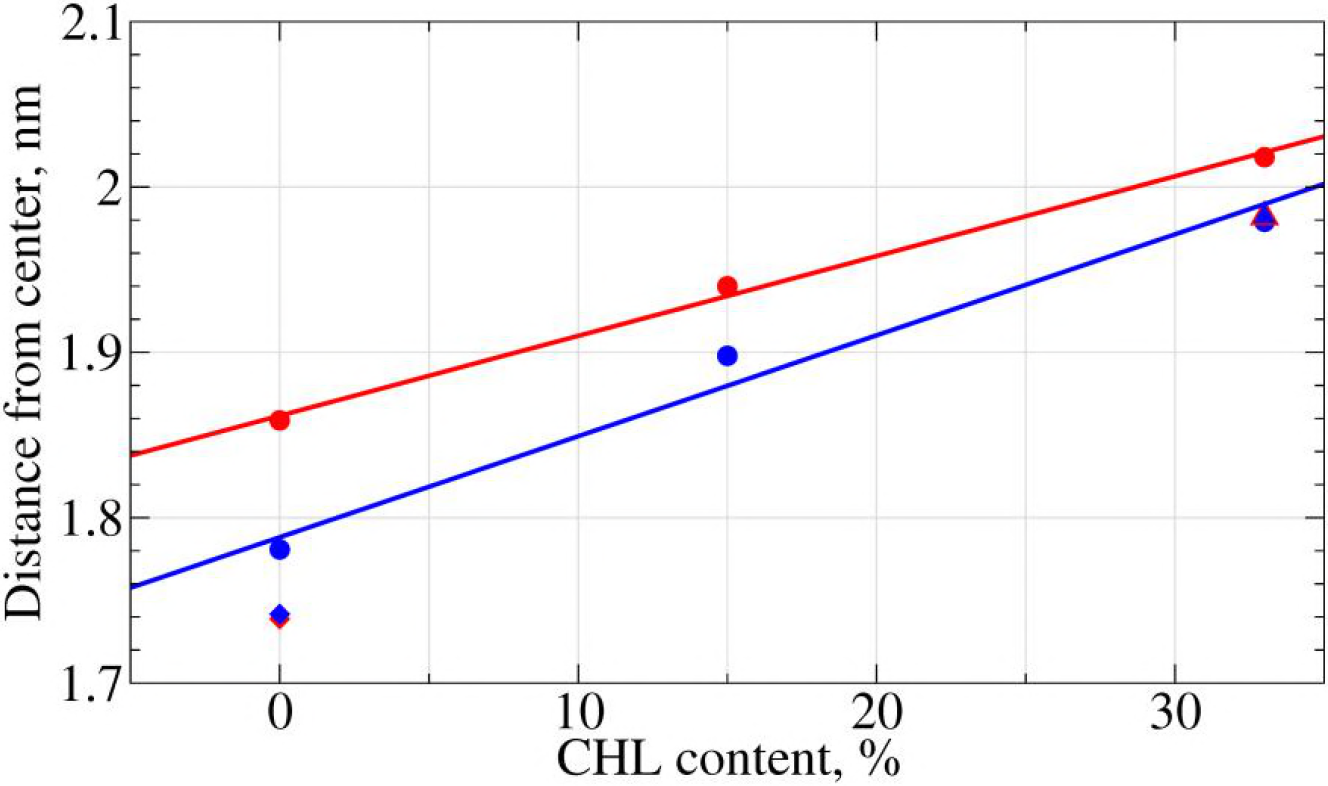
Absolute distance from center of the membrane to the peak of head groups density for different membrane models as a function of cholesterol content. The red and the blue lines are the linear regressions of the values for the normal model with different CHL content, for the inner and outer leaflets respectively. The red color refers to the inner leaflet and the blue to the outer leaflet. Circles refer to the normal membrane models, triangles to the cancer model and diamond to the DOPC model.

**Figure 4.**
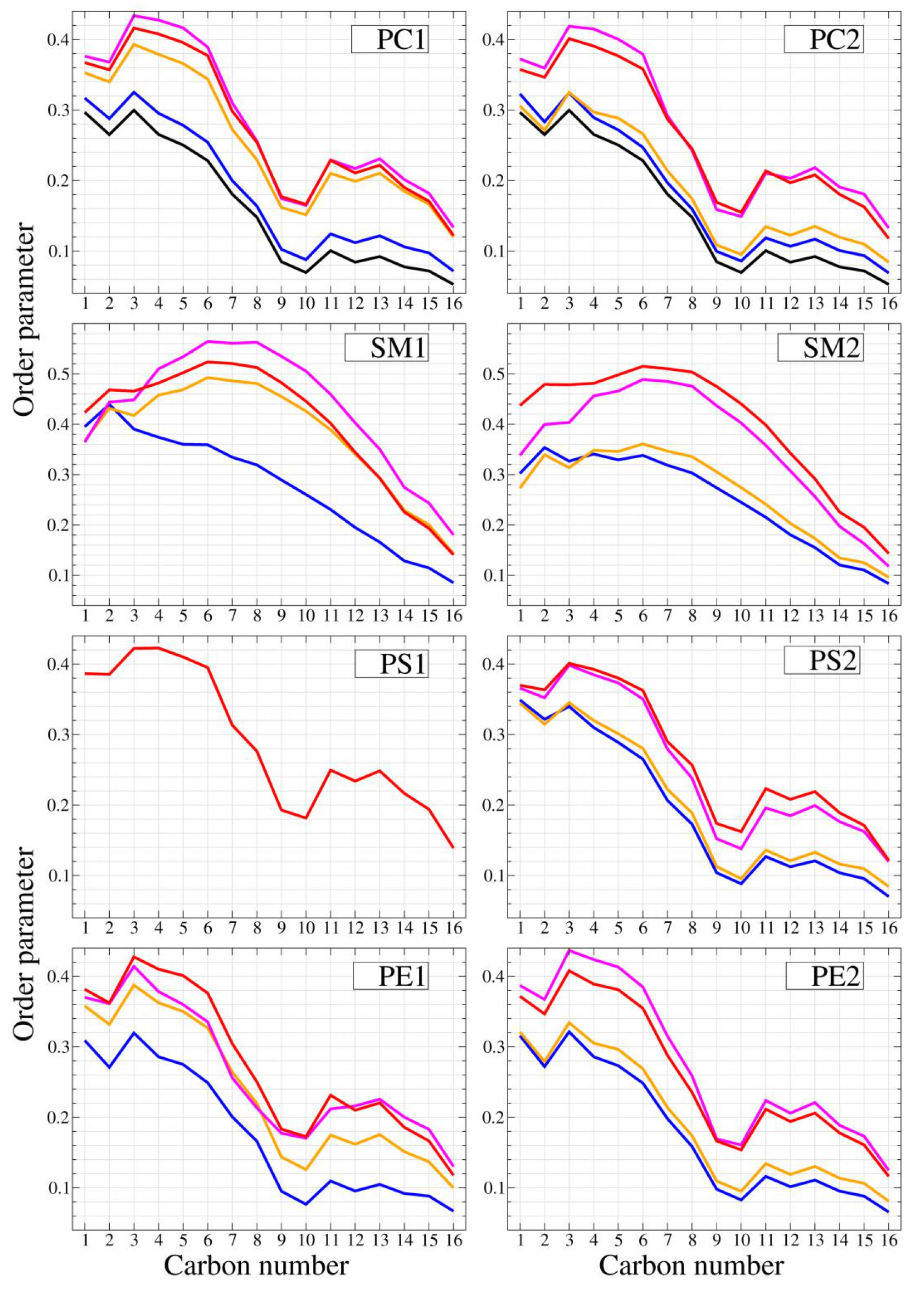
Order parameter of the lipid tails for all studied phospholipid species in the inner and outer monolayers for the different membrane models. Suffixes “1” and “2” correspond to outer and inner leaflets respectively.

The density profiles of cholesterol for normal and cancer membranes at 33% CHL content are comparable. For both normal and cancer membranes, the inner and outer monolayers contain approximately the same number of CHL molecules (the density profiles are symmetric). In our setup the CHL molecules could diffuse freely over the bicelle caps and redistribute between the monolayers. This may explain the visible asymmetry of the cholesterol distribution in the system with 15% of cholesterol (see Discussion for details).

Such redistribution of cholesterol influences the density of the lipid tails. It is evident from Fig. 2 that the density of the tails decreases substantially upon increase of cholesterol content. This is expectable since CHL molecules intercalate between the tails and increase the total volume of the monolayer. The redistribution of CHL to the outer monolayer correlates with smaller tails density in this monolayer at 15% cholesterol content.

The order parameters of the lipid tails also show remarkable dependence on cholesterol content. All phospholipids (PC, PE and PS) show very similar trends. The order parameter increases with the increase of cholesterol content in the corresponding monolayer. This general trend is logical and easily explainable by the fact that rigid CHL molecules intercalate between the lipid tails and increase their ordering. Such behavior has already been observed in pure 2,3dipalmetoyl-D-glycero-1-phosphatidylcholine (DPPC) and 3-palmetoyl-2-oleyl-D-glycero-1-phosphatidylchone (POPC) model membrane (15, 98). Asymmetric distribution of cholesterol at 15% is also clearly visible by different position of the corresponding curve for inner and outer monolayers (the orange curve is much higher for PC1 and PE1 than for PC2 and PE2).

The ordering of PC tails in pure DOPC membrane is lower than the ordering of PC tails in the normal membrane with 0% cholesterol. This is explained by the influence of more ordered saturated SM tails in the normal membrane.

The difference in order parameter between normal and cancer membrane with 33% of cholesterol are rather small. The tails of PC and PE lipids are less ordered in the cancer membrane while the tails of PS are more ordered in the inner monolayer. This is easily explainable by the redistribution of SM lipids. In the cancer membrane SM is distributed evenly between the monolayers, while in the normal membrane most of SM is in the outer leaflet. This leads to a decrease in ordering in the outer monolayer and its increase in the lower monolayer.

The ordering of the tails of SM lipids themselves follows the same pattern except remarkable difference of the curves with low content of cholesterol (SM1 at 0% and SM2 at 0% and 15%). In the absence of cholesterol the order of saturated SM tails decreases monotonously towards the center of the membrane. In contrast, in the presence of cholesterol the ordering increases and reaches the maximum in the middle of the tail near carbons 7 and 8. This clearly demonstrates formation of strong SM/CHL complexes. The similar changes of the order parameter were also reported in the case of pure DPPC membranes with an increasing cholesterol content (98) which is in line with well-known affinity of cholesterol to saturated lipid tails (11).

It is thus possible to infer that cholesterol concentration changes the physical properties of the membrane more than the redistribution of the lipids between the leaflets.

### Permeation of cisplatin

Figure 5A shows the PMFs of cisplatin permeation through the membranes under study. There is a high energy barrier in the center of the membrane, which is expected for such hydrophilic compound as cisplatin. The height of the barrier is the smallest for pure DOPC membrane (~40 kJ/mol) which is comparable to previous results obtained in a pure 1,2-dimyristoyl-sn-glycero-3-phosphocholine (DMPC) membrane (~50 kJ/mol) (99). For normal membrane with 0% cholesterol it is much higher (~62 kJ/mol) and increases even more with the increase of CHL content (up to 70 kJ/mol). The difference in the barrier height between the systems with 0% and 15% cholesterol is minimal, but its width is much larger for 15% system. It is remarkable that the width of the barrier increases mostly in the outer monolayer, which is enriched in cholesterol in the system with 15% CHL content. The PMFs for normal and cancer membranes at 33% CHL content are almost identical.

**Figure 5.**
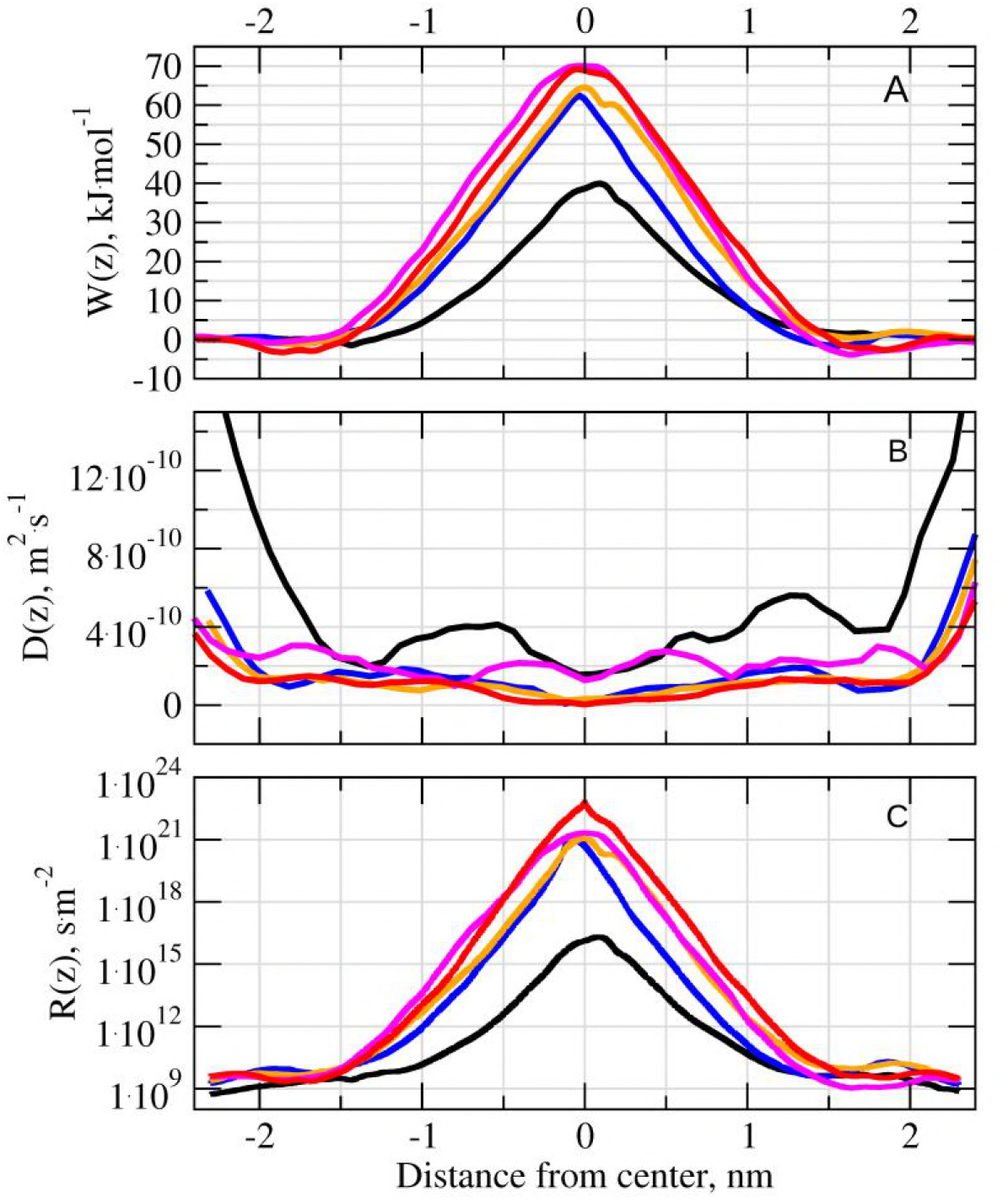
A: The top panel shows the PMF of cisplatin, W(z), B: the middle panel shows the local diffusion coefficient of cisplatin, D(z) and C: the bottom panel shows the local resistance of cisplatin R(z) for the different models of membranes. R(z) is represented in log scale.

Figure 5B shows diffusion coefficient of cisplatin in different systems. Again, pure DOPC membrane shows the highest values of D. There are no significant differences in D between normal membranes with 0%, 15% and 33% of CHL content, which suggests that cholesterol does not influence transversal diffusion of cisplatin. In contrast, there is a clear difference between normal and cancer membranes. Diffusion of cisplatin in the cancer membrane is surprisingly slower especially in the center of bilayer and in the regions of head groups.

The figure 5C shows the resistance R of different membranes to the permeation of cisplatin. The curves follow the shape of the PMFs but their heights are modulated by the difference in diffusion coefficients. The integration of these curves gives the permeabilities presented in Table 2.

**Table 2.**
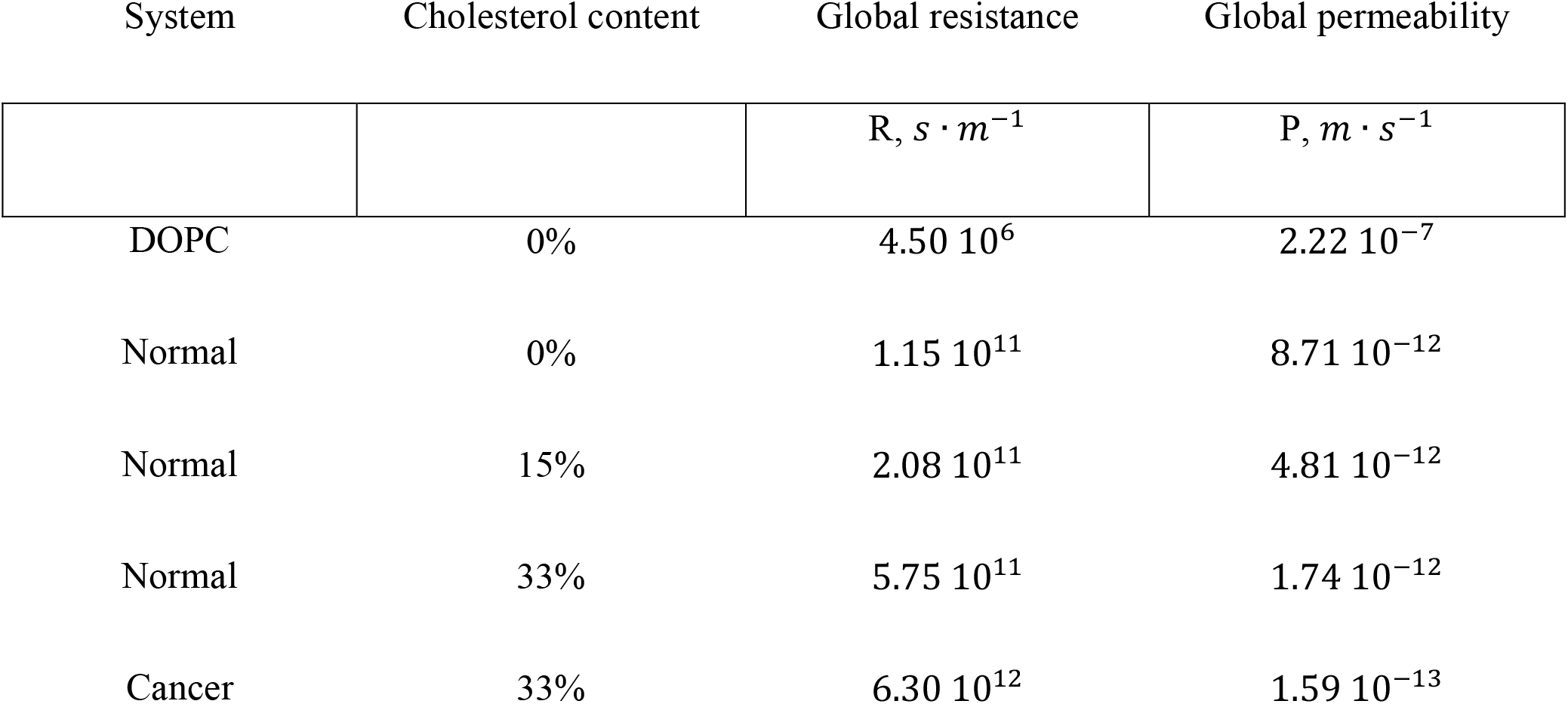
Values of the permeation resistances and permeabilities of cisplatin for different model membranes.

| System | Cholesterol content | Global resistance | Global permeability |
| --- | --- | --- | --- |
| | | $R, s \cdot m^{-1}$ | $P, m \cdot s^{-1}$ |
| DOPC | 0% | $4.50 \cdot 10^6$ | $2.22 \cdot 10^{-7}$ |
| Normal | 0% | $1.15 \cdot 10^{11}$ | $8.71 \cdot 10^{-12}$ |
| Normal | 15% | $2.08 \cdot 10^{11}$ | $4.81 \cdot 10^{-12}$ |
| Normal | 33% | $5.75 \cdot 10^{11}$ | $1.74 \cdot 10^{-12}$ |
| Cancer | 33% | $6.30 \cdot 10^{12}$ | $1.59 \cdot 10^{-13}$ |

It is clearly seen that the permeability is the largest for pure DOPC membrane and drops by 5 orders of magnitude in the model of plasma membrane with 0% cholesterol. Increase of cholesterol concentration from 0% to 33% decreases the permeability further by another order of magnitude. Finally, symmetrization of the lipid content of monolayer in the cancer membrane leads to even larger decrease of permeability. The cancer membrane appears to be ~11 times less permeable than the normal one.

## Discussion

### Composition of the membranes

Distribution of different lipid species in the membranes of cancer cells is usually discussed in terms of PS exposure, while the distribution of other lipids is rarely considered. This limits available experimental data concerning lipid content of the cancer cell membranes. Taking into account lack of reliable data we decided to consider the simplest model of the cancer cell membrane where distribution of all lipid species is the same in both monolayers. We are aware that this model is likely to be oversimplified and in reality the membrane is likely to remain somewhat asymmetric. However, fully symmetric model is the ideal reference system, which could be used to compare the membranes of different cancer cells if their lipid composition and distribution become available.

### The effect of cholesterol

The effect of cholesterol in our simulations is consistent with previous studies which suggest significant ordering and stiffening of the membrane upon increase of cholesterol concentration. Such stiffening and decrease of the flexibility of the lipid tails decreases the permeability of the membrane to cisplatin by approximately one order of magnitude in comparison with cholesterol-free membrane.

However, there is one puzzling observation in our results: a pronounced asymmetry of cholesterol density distribution in the system with 15% of cholesterol. The question of cholesterol distribution in large heterogeneous and asymmetric systems has only rarely been addressed in MD simulations, but uneven distributions of cholesterol between the leaflets of two model membranes has been reported recently by Ingólfsson et al. (4, 5). Close inspection of our system shows that the asymmetry comes from only three cholesterol molecules, which go from inner to outer monolayer in course of equilibration. Thus statistical significance of observed asymmetry is questionable since it could be easily caused by random motion of few cholesterol molecules rather than systematic reasons. At the same time there is a well-established strong affinity of cholesterol to sphingomyelin which may drive cholesterol molecules to SM-reach outer monolayer of the normal membrane. It is possible to speculate that cholesterol tends to populate the outer leaflet up to a saturation concentration first and then redistributes to the inner leaflet. 15% cholesterol content is likely to be insufficient to saturate the outer leaflet completely which may cause the observed asymmetry. In contrast, 33% cholesterol content is enough for saturation which leads to equal distribution between the monolayers. One should consider this speculation with care because lateral diffusion and the flip-flop processes should be sampled much better to draw reliable conclusion. It is rather straightforward to test this hypothesis in a series of simulations with different cholesterol distributions but this is beyond the scope of the current work.

### Permeability in comparison to experimental results

It is very important to compare our results with permeabilities of the membranes of model vesicles and living cells to cisplatin available in the literature. The majority of experimental works reports the kinetics of cisplatin permeation through the lipid membranes of cells and artificial liposomes as a single-step first order process (100). Thus, an effective kinetic constant of permeation, *k* is usually measured in such studies. A relation between permeability and effective kinetic constant is non-trivial in general case and depends on the membrane area, volume of the cell or liposome and membrane heterogeneity (101-103). Unfortunately there are no experimental works where the area to volume ratio of the cells is determined at the same time as the kinetic constant of cisplatin permeation. That is why in order to estimate the permeability from available kinetic data we have to use an oversimplified model of spherical cell with radius *r*, which was initially proposed for the liposomes (101-103):

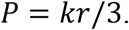

We considered three experimental works, which allow determining the permeabilities of cisplatin in either cells or artificial membranes (Table 3).

**Table 3.** Values of permeabilities for cisplatin estimated from the literature.

| Author | System | Permeability |
| --- | --- | --- |
| | | $P, m \cdot s^{-1}$ |
| Nierwzicki *et al.* (99) | DMPC bilayer (MD simulations) | $6.67 \cdot 10^{-7}$ |
| Eljack *et al.* (65) | DOPC vesicles<br>($[Cl^-] = 150 mM$) | $1.06 \cdot 10^{-8}$ |
| | DOPC vesicles<br>($[Cl^-] = 10 mM$) | $8.93 \cdot 10^{-10}$ |
| El-Kareh *et al.* (104) | A2780/CP3 human ovarian resistant cancer | $1.87 \cdot 10^{-9}$ |
| | COLO 205 colon carcinoma | $1.24 \cdot 10^{-9}$ |
| | A2780 human ovarian wild type cells | $1.21 \cdot 10^{-9}$ |
| | CAL 27 head and neck cancer | $1.19 \cdot 10^{-10}$ |
| | MKN45 gastric cancer | $1.01 \cdot 10^{-10}$ |
| | MKN74 gastric cancer | $4.90 \cdot 10^{-11}$ |
| | C32 human melanoma | $4.79 \cdot 10^{-11}$ |
| | G361 human melanoma | $2.21 \cdot 10^{-11}$ |
| Ghezzi *et al.* (68) | MCF-7 breast cancer | $6.57 \cdot 10^{-11}$ |

In the work of Eljack et al. (65) the permeation of cisplatin in pure DOPC vesicles was studied depending on the concentration of chloride ions in solution (the values for 10 mM and 150mM external Cl^-^ concentrations are shown in Table 3). Our results for pure DOPC membrane are one order of magnitude higher (~10^−7^ m/s in simulations versus ~10^−8^ m/s in experiment). It is necessary to note that only the counterions necessary to balance the system charge are present in our simulations. We also did not consider complex interplay of reactions between different cisplatin derivatives which is present in salt solutions and may influence an effective concentration of pristine cisplatin in experiment. Furthermore, the membranes of our systems are flat while the membranes of real cells and liposomes are curved significantly. We plan to investigate the influence of the curvature on the permeation of the drugs in our future studies but it is clear from multiple indirect evidence that the curvature should affect the permeability. That is why our setup is not directly comparable to experimental conditions which may explain observed discrepancies.

The work of Ghezzi *et al*. (68) reports a time-dependent uptake of cisplatin in breast cancer MCF-7 cell line. Considering the uptake of cisplatin as a first order kinetic process, we fitted the evolution of the intracellular concentration by an exponential function to obtain the kinetic constant (see supplementary information for the details). The average volume of the cells is estimated as 2 pL in this work. We assumed that the cells are spherical and computed the permeability. Obtained value of ~10^−11^ m/s is 1-2 orders of magnitude higher than our simulation results.

Finally, the work of El-Kareh *et al*. (104) reports the kinetic constant of cisplatin in different cell lines, which were computed in order to build a pharmacokinetic model for several platinum(II) species. Since the volumes and the area to volume ratios of the studied cells are not reported in this paper we assumed that the cells were spherical and that their volume was the same (2 pL) as in the work of Ghezzi *et al*. (68) mentioned above. This allowed us to obtain a rough estimate of permeabilities. However, obtained values are likely to be underestimated since the cell are not spherical and their real area to volume ratio will tend to be higher since real system exhibits complex shape with high membrane curvature.

The only computational work which is directly comparable with the present study is the paper of Nierwzicki *et al*. (99) where the permeability of cisplatin in a pure DMPC bilayer was reported as 6.67 10^−7^*m* · s^−1^. This value is of the same order of magnitude as our result for pure DOPC bilayer.

It is possible to conclude that direct quantitative comparison between MD simulations of permeability and the data obtained experimentally on real cell lines remains challenging. The largest uncertainity comes from the fact that area to volume ratio of the studied cells is not estimated in experimental works. Without these data one is restricted to the primitive and inaccurate model of the cell as spherical homogeneous vesicle, which can easily lead to severe underestimation of the permeability by several orders of magnitude. Taking this into account our results look reasonable and consistent.

Although our simulation setup is not directly comparable to experimental conditions it is much closer to real membranes than the vast majority of membrane models routinely used in MD studies nowadays. We demonstrated that taking into account realistic lipid composition and cholesterol content brings estimated permeabilities much closer to experimental values in comparison to simplified membrane models. We expect that accounting for membrane curvature in future works will reproduce experimental results even better. In general we believe that MD simulations of realistic curved and asymmetric membranes with variable cholesterol content will soon allow for reliable estimation of permeabilities for various compounds including cisplatin and other platinum drugs.

## Conclusion

In this work we studied the influence of lipid composition and cholesterol content on the permeation of cisplatin through the model membranes of normal and cancer cells.

It is shown that the loss of lipid asymmetry in the cancer membranes leads to decrease their permeability to cisplatin by one order of magnitude in comparison to the asymmetric membranes of normal cells. This effect is caused by slower diffusion of cisplatin in the cancer membrane while the energy barrier of permeation remains the same. It is possible to speculate that this effect may contribute to cisplatin resistance in the cancer cells, which exhibit pronounced loss of lipid asymmetry.

The change of cholesterol molar ratio from 0% to 33% also decreases the permeability of the membrane by approximately one order of magnitude. Thus, changes of cholesterol content in cancer cell membranes may influence their permeability to cisplatin significantly.

It is also shown that the single-component DOPC membrane is a very poor model for cisplatin permeation in real cells since its permeability is 5-6 orders of magnitude higher than one of the membrane with realistic lipid composition.

We also conclude that direct quantitative comparison between MD simulations of permeability and the data obtained experimentally on real cell lines remains challenging due to the lack of models, which take into account complex shape of the cells, and uncertainty related to chemical modifications of cisplatin in solution.

## Acknowledgments

This work was supported by the European Union’s Horizon 2020 research and innovation programme under the Marie Skłodowska-Curie grant agreement No. 690853. The computational studies work was performed using HPC resources from GENCI-[TGCC/CINES/IDRIS] (Grant 2015-[c2016077586]) and the Centre de calcul regional Romeo. We would like to address special acknowlegments to the Mésocentre de calcul de Franche-Comté for their important support in this work.

The authors declare no competing financial interest.

## Authors contributions

CR and SY designed the study. SY developed simulation methodology. SY and TR developed software for data analysis. TR performed simulations, analyzed and visualized data, performed comparison with experimental results. CR guided discussion and interpretation of the results. The manuscript was written by all authors.

